# Omicron BA.1 and BA.2 Variants Increase the Interactions of SARS-CoV-2 Spike Glycoprotein with ACE2

**DOI:** 10.1101/2021.12.06.471377

**Authors:** Mert Golcuk, Ahmet Yildiz, Mert Gur

## Abstract

SARS-CoV-2 infection is initiated by binding of the receptor-binding domain (RBD) of its spike glycoprotein to the peptidase domain (PD) of angiotensin-converting enzyme 2 (ACE2) receptors in host cells. Recently detected Omicron variant of SARS-CoV-2 (B.1.1.529) is heavily mutated on RBD. Currently, the most common Omicron variants are the original BA.1 Omicron strain and the BA.2 variant, which became more prevalent since it first appeared. To investigate how these mutations affect RBD-PD interactions, we performed all-atom molecular dynamics simulations of the BA.1 and BA.2 RBD-PD in the presence of full-length glycans, explicit water and ions. Simulations revealed that RBDs of BA.1 and BA.2 variants exhibit a more dispersed interaction network and make an increased number of salt bridges and hydrophobic interactions with PD compared to wild-type RBD. Although BA.1 and BA.2 differ in two residues at the RBD-ACE2 interface, no major difference in RBD-PD interactions and binding strengths were observed between these variants. Using the conformations sampled in each trajectory, the Molecular Mechanics Poisson-Boltzmann Surface Area (MMPBSA) method estimated ~34% and ~51% stronger binding free energies for BA.1 and BA.2 RBD, respectively, than wild-type RBD, which may result in higher binding efficiency of the Omicron variant to infect host cells.

## Introduction

The recent appearance and the rapid rate of infection of a heavily mutated B.1.1.529 variant of SARS-CoV-2, named Omicron, have raised concerns around the world, with many countries temporarily limiting their international travel. World Health Organization has designated the omicron variant as a variant of concern (VOC)^1^. Currently, the omicron variant has three major sub-lineages, namely BA.1, BA.2, and BA.3.^2^ BA.1 became the first dominant omicron and together with BA.2 they are currently the most observed SARS-CoV-2 variants. The BA.2 variant accounts for half of all new SARS-CoV-2 cases globally.^3^ The BA.1 variant comprises 30 mutations on the spike glycoprotein (S), while the BA.2 variant comprises 28. Remarkably, 15 and 16 of these mutations are located on the receptor-binding domain (RBD) of the BA.1 and BA.2 variants, respectively. Among these RBD mutations 12 (G339D, S373P, S375F, K417N, N440K, S477N, T478K, E484A, Q493R, Q498R, N501Y, and Y505H) are shared among the BA.1 and BA.2 variants (Figure 1).

**Figure 1.**
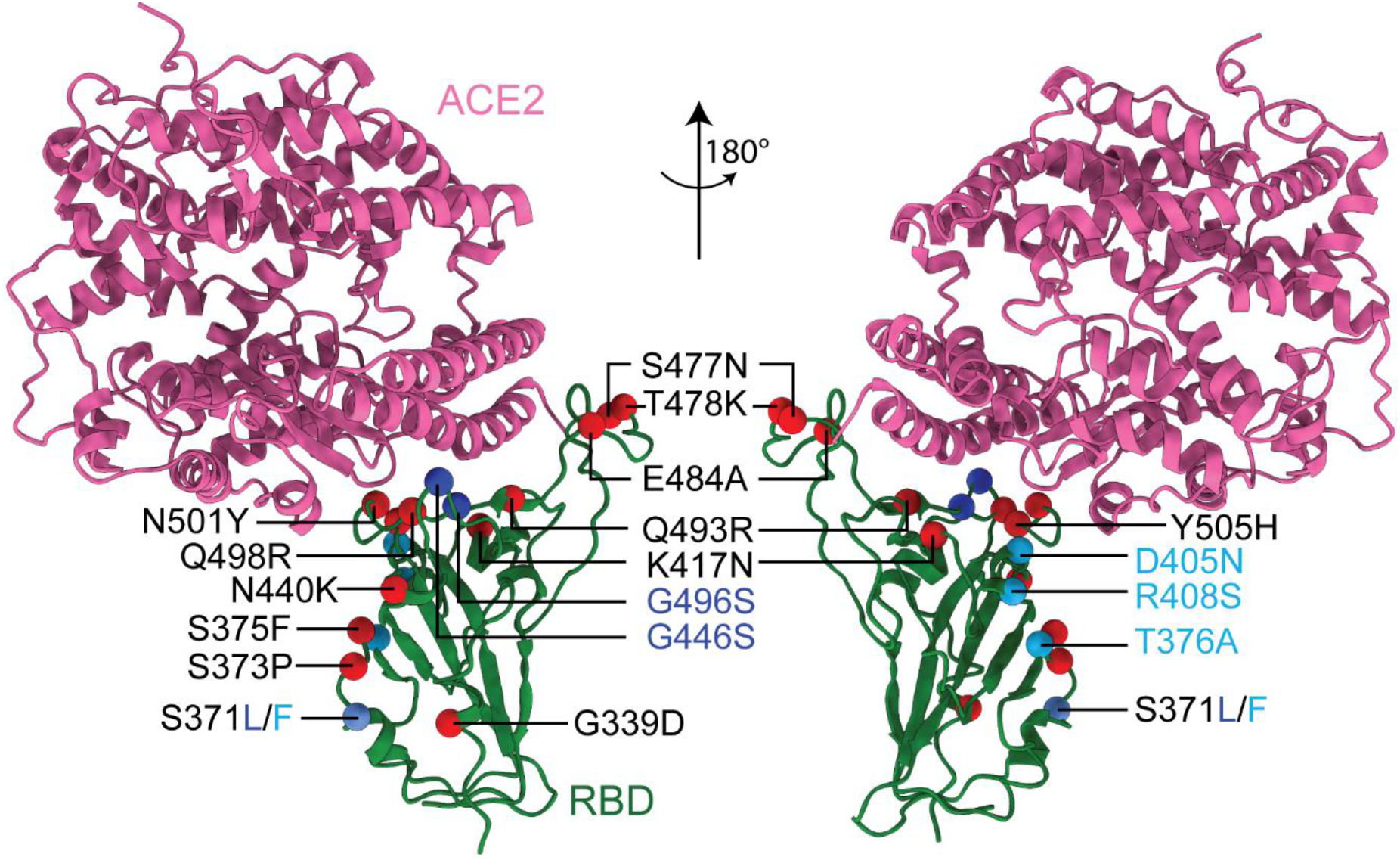
Location of RBD mutations for the Omicron variant. Mutations found on both BA.1 and BA.2 are highlighted with red beads, while the mutations specific to BA.1 and BA.2 variants are highlighted with blue and turquoise colored beads, respectively.

RBD interacts with the peptidase domain (PD) of angiotensin-converting enzyme 2 (ACE2) receptors and plays a critical role in the host cell entry of the virus. RBD is a critical antibody and drug target, and all the available vaccines produce antibodies that neutralize the RBD-PD interaction. Mutations on both BA.1 RBD (RBD_BA.1_) and BA.2 RBD (RBD_BA.2_) are surface-exposed and being targeted by various antibodies (Figure S1) and nanobodies. In addition, for BA.1, 11 of these 15 mutations are located on the ACE2 binding interface, while for BA.2 nine of these are located on the ACE2 binding interface (Figure 1). For both BA.1 and BA.2 four hydrophilic residues mutated to positively charged residues (N440K, T478K, Q493R, and Q498R), one negatively charged residue mutated to hydrophobic residue (E484A), one positively charged residue mutated to hydrophilic residue (K417N), and for hydrophilic residues are mutated to again hydrophilic residues (S477N, N501Y, and Y505H) at RBD’s PD binding interface. In addition, to these mutations, two neutral residues mutated to hydrophilic residues (G446S and G496S) in BA.1. Thus, both RBD_BA.1_’s and RBD_BA.2_’s PD binding interface are more positively charged than RBD_WT_. Furthermore, the PD binding interface of RBD_BA.1_ comprises more hydrophilic residues than RBD_BA.2._ Our previous all-atom Molecular Dynamics (MD) simulations^4^ showed that 5 of these mutated residues form pairwise interactions between wild-type (WT) S and ACE2 (salt bridges between K417-D30 and E484-K31, and hydrogen bonding between Q493-E35, Q498-Q42, Q498-K353, and Y505-E37). It is still unclear how BA.2 omicron mutations affect the binding strength of RBD to ACE2 and the ability of existing SARS-CoV-2 antibodies to neutralize this interaction. Furthermore, the difference in binding characteristics and strength of omicron BA.1 and BA.2 variants remains to be explored.

In order to explore the effect of various omicron variant mutations on RBD-ACE2 interactions, we performed an extensive set of MD simulations of the RBD-PD complex for the two main omicron variants BA.1 and BA.2. Our simulations totaling 3 μs in length revealed that both RBD_BA.1_ and RBD_BA.2_ exhibit a more dispersed interaction network on the RBD-ACE2 interaction surface compared to WT RBD (RBD_WT_). Furthermore, an increased number of salt bridges and hydrophobic interactions of RBD_BA.1_ and RBD_BA.2_ with PD were observed. Molecular Mechanics Poisson-Boltzmann Surface Area (MMPBSA) method estimated ~34% and ~51% stronger binding free energy for RBD_BA.1_ and RBD_BA.2_, respectively, compared to RBD_WT_.

## Results

### RBD_BA.1_-PD Interactions

We performed all-atom MD simulations of the RBD_BA.1_-PD in the presence of full-length glycans on both S RBD and ACE2 PD, ^5,6^ explicit water and ions (~200k atoms in total). Four sets of MD simulations each of 300 ns in length were performed using the parameters of our previous RBD-PD simulations for the WT,^4^ alpha, and beta variants.^7^ These four sets of simulations were combined into a single 1,200 ns long trajectory to investigate the RBD_BA.1_-PD interactions. Simulations revealed a more extensive interaction network for RBD_BA.1_-PD with PD compared to RBD_WT_. We detected five salt bridges between RBD_BA.1_ and PD; one of them (K440-E329) medium and four (R403-E37, R493-E35, R493-D38, and R498-D38) with high frequency (Figure 2). In comparison only 2 high frequency salt bridges existed between RBD_WT_ and PD^4^ (Figure 2 and Table S1) and both of those disappeared in the BA.1 variant (Figure 2 and Table S2). The RBD_BA.1_ forms all of the ten high frequency hydrophobic interactions that were observed for RBD_WT_-PD and an additional high frequency hydrophobic interaction between Y501-Y41. Compared to eight hydrogen binding between RBD_WT_ and PD (3 high and five medium frequency), six hydrogen bonds were observed between RBD_BA.1_ and PD (3 high and 3 medium frequency). Only two of these interactions were also observed for the WT, while other four are newly formed (Figure 2). Collectively, the total number of salt bridges, hydrophobic interactions, hydrogen bonds at the S-ACE2 interface changed by 150%, 10% and −25%, respectively.

**Figure 2.**
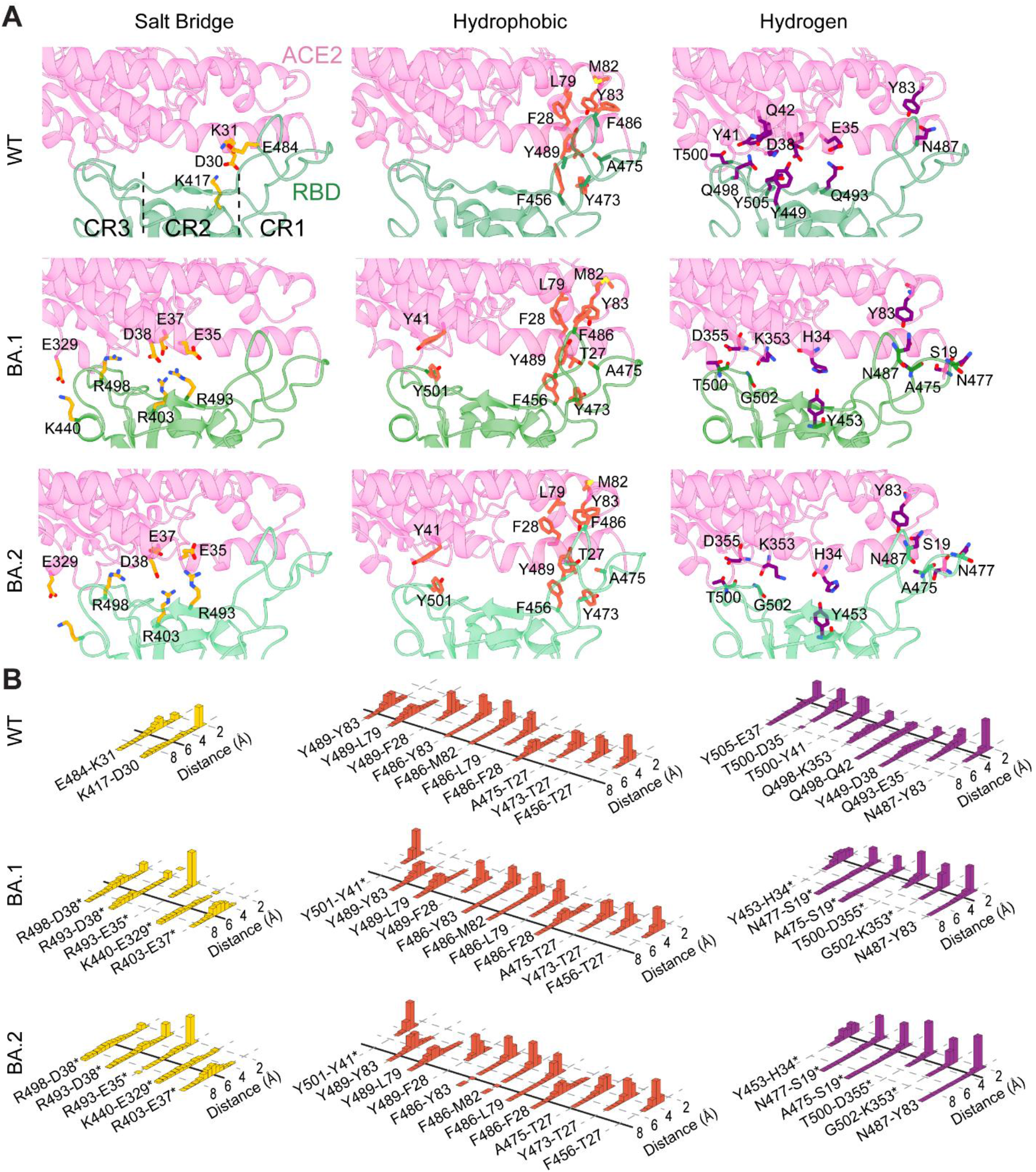
(A) Interactions between RBD_WT_, RBD_BA.1_ and RBD_BA.2_ of the SARS-CoV-2 S protein and the PD of human ACE2. Representative snapshots of the all-atom MD simulations highlight salt bridges, hydrophobic interactions, and hydrogen bonding between RBD_WT_-PD, RBD_BA.1_-PD and RBD_BA.2_-PD. The interaction surface is divided into three distinct regions (CR1-3)^4,12^ (B) Normalized distributions of the distances between the amino-acid pairs that form salt bridges (orange), hydrophobic interactions (red), and hydrogen bonds (purple) between RBD_WT_, RBD_BA.1_ and RBD_BA.2_ and PD. Newly formed interactions due to mutations are marked with an asterisk. Solid lines represent the minimal threshold distance between these residues to form each class of pairwise interactions.

Our simulations also revealed a change in the spatial distribution of RBD-PD interactions along the interaction surface due to the mutations in the BA.1 variant, which are mostly consistent with recently reported RBD_BA.1_-ACE2 structures.^8–11^ Between RBD_WT_ and PD, salt bridges are concentrated at the interface of contact region 1 (CR1) and CR2, while hydrogen bonding and hydrophobic interactions are concentrated in CR3 and CR1, respectively (Figure 2A).^4^ In comparison, RBD_BA.1_ exhibits a more dispersed interaction network along the RBD-PD interaction surface (Figure 2). RBD_BA.1_ mutations result in 2 additional interactions (hydrogen bonds) in CR1, 4 additional interactions (3 salt bridges and 1 hydrogen bond) in CR2, and 5 additional interactions (2 salt bridges, 2 hydrogen bonds, and 1 hydrophobic interaction) in CR3. Furthermore, RBD_BA.1_ mutations result in the loss of 1 interaction (1 salt bridge) in CR1, 3 interactions (1 salt bridge and 2 hydrogen bonds) in CR2, and 5 interactions (5 hydrogen bonds) in CR3. This may result in an altered binding mechanism and negatively impact the current inhibition mechanism by neutralizing antibodies and nanobodies.

### RBD_BA.2_-PD Interactions

For RBD_BA.2_-PD, four sets of all-atom simulations each of 300 ns in length were performed in the presence of explicit water and ions. These simulations were combined into a single MD trajectory of 1,200 ns in length to investigate RBD_BA.2_-PD Interactions. As was the case for RBD_BA.1_, RBD_BA.2_ showed a more extensive interaction network with PD compared to RBD_WT_. Between RBD_BA.2_ and PD a total of five salt-bridges (two high and three medium frequency), 11 hydrophobic interactions (all high frequency), and six hydrogen bonding (five high and one medium frequency) (Figure 2 and Table S2), which corresponds to a 150%, 10% and −25% changed compared to those observed for RBD_WT_-PD, respectively. Similar to BA.1, RBD_BA.2_ exhibits a dispersed interaction network for each type of interaction type along the RBD-PD interaction surface (Figure 2).

The difference in the RBD-PD interface for BA.1 and BA.2 variants are the two residues located at residue positions 446 and 496, which are S446 and S496 for BA.1 and G446 and G496 for BA.2. Yet, the interacting RBD-PD residue pairs for BA.1 and BA.2 were identical. Comparing the frequencies of RBD_BA.2_-PD interactions with RBD_BA.1_-PD shows that the total number of high frequency hydrogen bonds increased by three, while high frequency salt bridges decreased by two for BA.2 with respect to BA.1. Conclusively, the binding interactions network for RBD_BA.1_–PD and RBD_BA.2_–PD share similar features, differing only in the observation frequencies in 6 out of 22 interactions.

### Effect of omicron BA.1 and BA.2 mutations on RBD, PD, and their interface fluctuations

To investigate the effect of omicron mutations onto the RBD binding dynamics, we quantified the Root Mean Square Fluctuations (RMSF) of the C_α_ atoms of the RBD residues located on the PD binding surface for the RBD-PD compelxes.^4^ All sets of MD simulations performed for WT, BA.1, and BA.2 were combined into a single trajectories of 600 ns, 1,200ns, and 1,200 ns lengths, respectively. The rigid body motions were eliminated for each trajectory by aligning the RBD interacting surface of PD for each conformer with their starting crystal structure. Both BA.1 and BA.2 mutants of RBD had lower residue fluctuations at the interface suggesting that tighter and more rigid binding compared to WT (Figure 3). At CR1 and CR3, BA.2 mutations caused a significantly larger decrease in fluctuations compared to BA.1, while in CR2 the opposite was observed (Figure 3).

**Figure 3.**
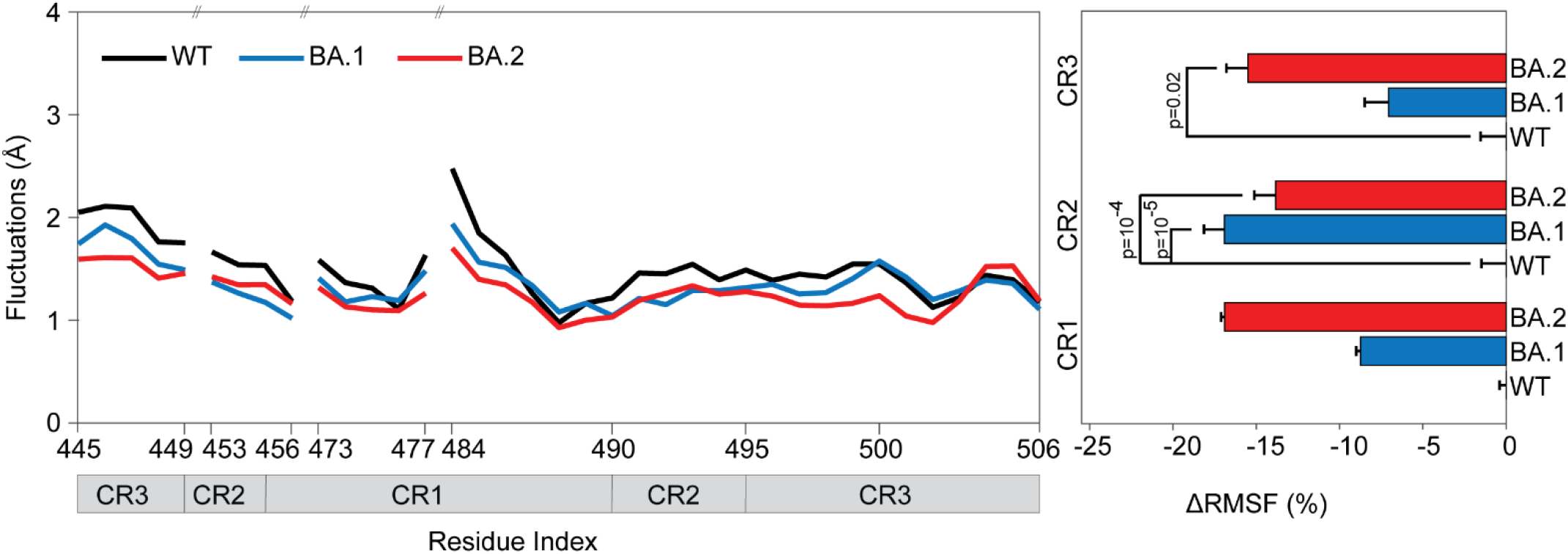
(Left) Effect of omicron mutations onto RBD and PD fluctuations. (Right) RMSF of RBD residues located on the PD binding surface of WT, BA.1 and BA.2 variants. P values were calculated using a two-tailed Student’s t-test. P values larger than 0.05 are not shown. Error bars represent standard deviation (s.d.).

We also aligned each trajectory with the RBD_WT_–PD crystal structure by the RBD beta sheet and helix C_α_ atoms. The RMSD of the beta sheet and helical regions of the RBD_BA.1_ and RBD_BA.2_ to RBD_WT_ resulted in trajectory average values of 0.82 Å and 0.79 Å, respectively (Figure S2). BA.1 and BA.2 mutations did not affect the RBD structure significantly, showing practically identical structures with WT.

### Binding free energies for RBD_BA.1_-PD and RBD_BA.2_-PD

Binding free energies of two sets of RBD_WT_-PD, four sets of RBD_BA.1_-PD, and four sets of RBD_BA.2_-PD simulations were calculated via the MMPBSA method^13,14^ using the VMD^15^ plugin CaFE^16^ (Table S3). MMPBSA calculations estimated 34% stronger binding free energy (−40 ± 9.7 kcal/mol, mean ± s.d., N = 4 sets) for RBD_BA.1_ compared to RBD_WT_ (−29.9 ± 7.3 kcal/mol, N = 2 sets). For RBD_BA.2_ MMPBSA calculations estimated 51% stronger binding free energy (−45.3 ± 9.1 kcal/mol, N = 4 sets) compared to RBD_WT_ (Figure 4). Considering that BA.1 and BA.2 induced small changes in the total percentage in the number of hydrophobic interactions and hydrogen bonds, while their effect on the total number of salt bridges was considerably large, we conclude that the increase in the number of salt bridges in the S-ACE2 interface resulted in this higher binding strength of RBD_BA.1_ and RBD_BA.2_ to PD, which may result in a higher efficiency of the SARS-CoV-2 virus to infect host cells.

**Figure 4.**
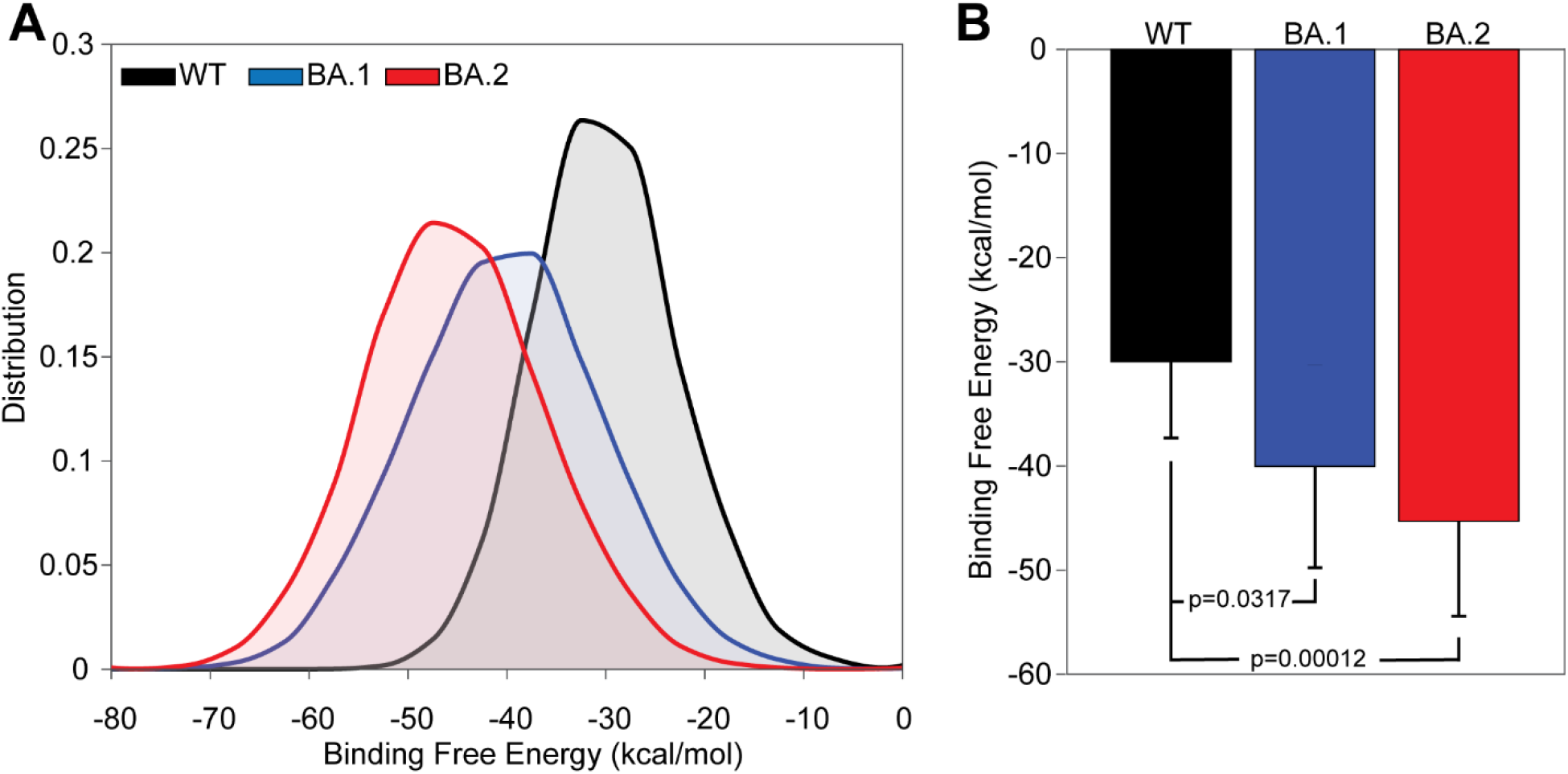
Binding free energies of RBDs to PD. (A) Distribution of the binding free energies of RBD_WT_, RBD_BA.1_, and RBD_BA.2_. (B) Mean binding free energy values of the RBD_WT_, RBD_BA.1_, and RBD_BA.2_ to PD. Error bars represent s.d. P values were calculated using a two-tailed Student’s t-test.

## Discussion

An extensive set of MD simulations totaling 3 μs in length were performed to investigate the effect of the two most common omicron variants BA.1 and BA.2 in RBD-PD interactions. The preprint of this study was the first in the literature to show via all-atom MD simulations the effect of the omicron BA.1 mutations on RBD-PD interactions and binding strength. Our findings have been supported with recent computational^17,18^ and experimental studies^11,19-21^. There is currently no consensus regarding the exact binding energy of the BA.1 omicron variant S protein to ACE2. Binding free energies ranging from −107.04 to −635.32 kcal/mol were reported by post-processing all-atom MD trajectories.^17,18^ Furthermore, K_D_ values ranging 25.3-38.9 nM were reported for the BA.1 omicron variant.^11,19–21^ While these approaches report different K_D_ values, consistent with our findings, they all estimate a higher binding strengths for RBD_BA.1_ compared to the RBD_WT_. Yet, effect of BA.2 mutations on ACE2 binding have not yet been reported in the literature. The binding free energies we estimated via MMPBSA method exhibits a 34% and 51% increase in binding strength for RBD_BA.1_ and RBD_BA.2_ compared to RBD_WT_. The analysis of the pairwise interactions between RBD and PD provided a detailed insight into this increased binding strength of the omicron variants. The most striking change induced by BA.1 and BA.2 mutations was the net change of 3 additional salt bridges. Both RBD_BA.1_ and RBD_BA.2_ mutations were shown to decrease the fluctuation of RBD residues at the ACE2 binding interface. Collectively our result highlight that both omicron variants result in a more extensive interaction network, and a stronger and tighter binding. Our MD simulations also revealed differences in observation frequencies, which result in a stronger and tighter binding for BA.2 compared to BA.1.

RBD_BA.1_ and RBD_BA.2_ mutations may also affect the binding affinity and neutralizing capability of SARS-CoV-2 S antibodies and nanobodies. Neutralizing antibodies were categorized into 4 classes according to their binding regions and mechanisms.^22^ Mutations introduced by both BA.1 and BA.2 variants overlap with critical interactions for all of these antibody classes. For example, we expect E484A mutation to eliminate E484-R52 salt bridge and E484-S57 hydrogen bonds in H11-H4 and H11-D4 nanobodies, and E484-N56, and E484-Y335 hydrogen bonds in Ty1 nanobody. ^7,23,24^ Additionally, Q493R mutation would eliminate the hydrogen bonds Q493-Y104 and Q493-S104 in H11-H4, and H11-D4, respectively. Furthermore, mutations are expected to eliminate salt bridge E484-R96 and hydrogen bonds E484-H35, Q493-R97, and Q493-S99 between RBD and a class 2 antibody C002 (Figure 5). Consistent with this view, point mutation Q493R mutations was reported to decrease the RBD dissociation constant of C002 from 11 nM to 596 nM.^22,25,26^

**Figure 5.**
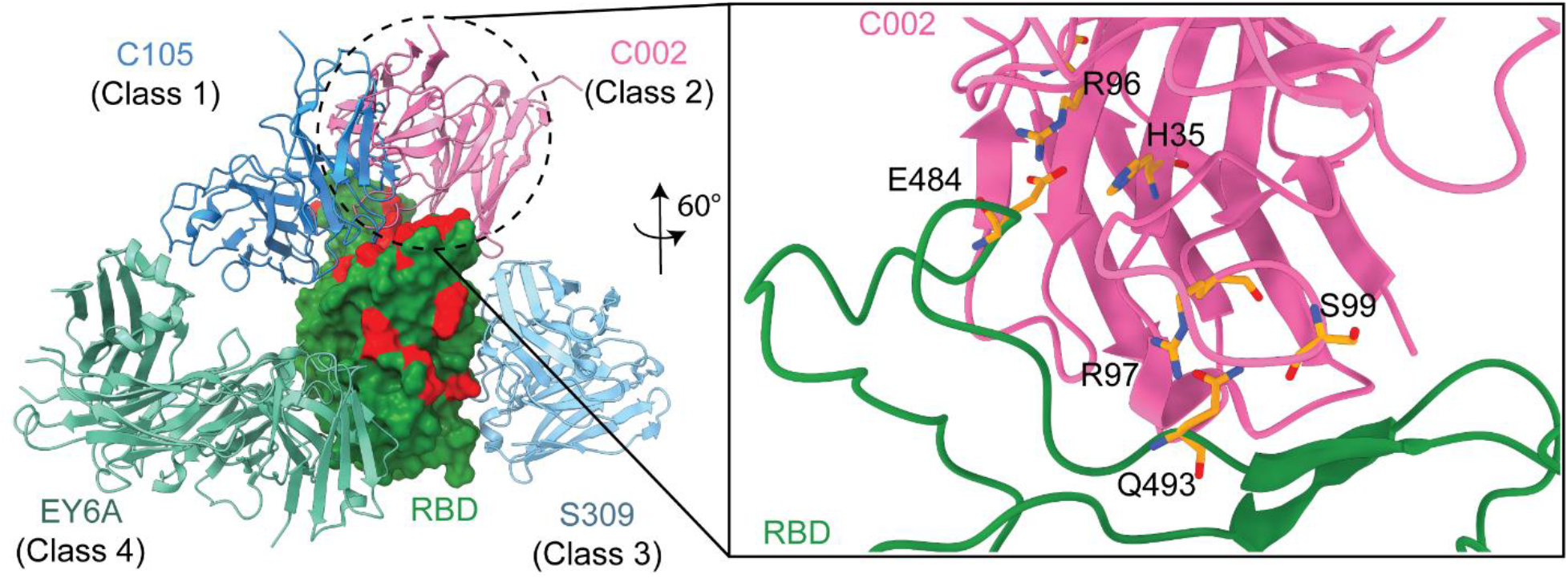
Class 1-4 Antibody binding poses on RBD. (Left) Binding poses of antibodies C105, C002, S309 and EY6A categorized as class 1-4 antibodies, respectively, are shown. Locations of RBD mutations of the omicron variant are shown in red. (Right) RBD-C002 interactions that are expected to be eliminated due to mutations in RBD_Omicron_ are shown on the right.

## Methods

### MD simulations system preparation

Systems were prepared in VMD^15^ as we previously performed RBD_WT_-PD, RBD_Alpha_-PD and RBD_Beta_-PD MD simulations.^4,7^ The RBD_BA.1_-PD structure^19^ did not exist when the preprint of this study was published. Thus, the structure of SARS-CoV-2 S protein RBD bound with ACE2 (PDB ID: 6M0J^27^) was used as a starting structure for the MD simulations of the RBD_BA.1_-PD complex. Omicron BA.1 variant RBD structure was modelled by introducing the 15 mutations located at the RBD of the SARS-CoV-2 omicron variant using the Mutator plugin of VMD^15^ onto the WT RBD structure. Chloride ion, zinc ion, and water molecules in the structures were kept. Since full length glycans are not visible in the crystal structure, we used glycan models.^5^ RBD_BA.1_-PD was solvated into a water box with 25 Å cushion in each direction using TIP3P model water molecules. Ions were added to neutralize the system and set the NaCl concentration to 150 mM.

The RBD_BA.1_-PD (PDB ID: 7T9L^19^) structure was published recently. Using this structure an additional solvated RBD_BA.1_-PD system was constructed. For the RBD_BA.2_-PD simulations, solvated RBD_BA.2_-PD systems were modeled by introducing L371F, T376A, D405N, and R408S mutations and reversing G446S and G496S Mutations in the RBD_BA.1_-PD structure, and subsequent solvating it and adding ions to the system.

### MD simulations

Two sets of conventional MD simulations each of 300 ns length (MD 1-2) were performed for the RBD_WT_-PD complex in the presence of explicit water molecules, ions and also full length glycans. Four sets of conventional MD simulations each of 300 ns length (MD 3-6) were performed for the RBD_BA.1_-PD complex in the presence of explicit water molecules, ions and also full length glycans. Three of those simulations were initiated RBD_WT_-PD based RBD_BA.1_-PD model (MD3-5), while one was initiated from the RBD_BA.1_-PD structure (MD6). Four sets of simulations each of 300ns length were performed for the RBD_BA.2_-PD complex in the presence of explicit water molecules, ions and also full length glycans (MD7-10). Prior to these production simulations, each system was minimized for 10,000 steps and then equilibrated for 2 ns by keeping the protein fixed. Subsequently, system was minimized for an additional 10,000 steps without fixing the protein, which is followed by 4 ns of equilibration with harmonic constraints applied on C_α_ atoms. All constraints were removed from the system and an additional 4 ns of MD simulations were performed; finalizing the minimization and equilibration steps prior to production runs.

MD simulations were performed in NAMD-2.14^28^, for MMPBSA calculations and system minimizations and equilibrations, and NAMD3^28^ for all production simulations under N, P, T conditions. CHARMM36^29^ force field and a time step of 2 fs was used in the simulations. Pressure was kept at 1 atm using the Langevin Nosé-Hoover method with an oscillation period of 100 fs and a damping time scale of 50 fs. Temperature was maintained at 310 K using Langevin dynamics with a damping coefficient of 1 ps^-1^. Periodic boundary conditions were applied in simulations and Particle-mesh Ewald method was used for long-range electrostatic interactions. 12 Å cutoff distance was used for van der Waals interactions.

### Criteria for Interaction Analysis

To determine salt bridge formation in MD simulations, a cutoff distance of 6 Å between the basic nitrogen and acidic oxygen was used.^30^, while for hydrophobic interactions, a cutoff distance of 8 Å between the side chain carbon atoms was used.^31–33^ A cutoff distance of 3.5 Å between hydrogen bond donor and acceptor, and a 30° angle between the hydrogen atom, the donor heavy atom and the acceptor heavy atom was used to determine hydrogen bond formation.^34^ Among those interaction pairs that satisfied the hydrogen bonding distance criterion but did not satisfy the angle criterion, were classified as electrostatic interactions. As was performed in our previous studies,^4,35^ observation frequencies of interactions sampled from MD simulations were classified as high and moderate for interactions that occur in 49% and above and between 15 and 48% of the total trajectory, respectively. Pairwise interactions with observation frequencies below 15% were excluded from further analysis.

### Binding Free Energy Predictions via MMPBSA method

For each set of simulation, 3000 snapshots each separated by 0.1 ns were selected from the simulations. The binding free energies were predicted for the RBD-PD complexes using the MMPBSA method^13,14^ which was conducted via VMD^15^ plugin CaFE^16^ as described by Liu and Tingjun.^16^ Entropy change during binding was neglected in calculations, consistent with previous MMPBSA calculations for RBD-PD interactions.^36,37^ Default parameters were used in CaFE^16^ calculations.

## Acknowledgements

This work is supported by COVID-19 HPC Consortium (Grant number: TG-BIO200053 and TG-BIO210181)

## Author contributions statement

Mert Gur and A.Y.,. Mert Gur supervised the project. Mert Golcuk and Mert Gur performed MD simulations. Mert Golcuk, A.Y., and Mert Gur prepared the manuscript.

