## Supporting Information for "Omicron BA.1 and BA.2 Variants Increase the Interactions of SARS-CoV-2 Spike Glycoprotein with ACE2"

**Effect of omicron BA.1 and BA.2 mutations on RBD and PD structures**

From the RBD distributions representative RBD conformers showing the most probable RMSD values were selected. Using these conformers the RMSD between the following pairs $RBD_{WT}$:$RBD_{BA.1}$, $RBD_{WT}$:$RBD_{BA.2}$ and $RBD_{BA.1}$:$RBD_{BA.2}$ were evaluated as 0.75 Å, 0.82 Å, and 0.75 Å, hence showing minimal difference.

**Table S1.** Pairwise interactions of $RBD_{WT}$ with ACE2.

| | **Interaction** | **Location** | **$RBD_{WT}$-ACE2**[2] |
|---|---|---|---|
| **Salt bridges** | K417-D30 | CR2 | 99% |
| | E484-K31 | CR1 | 80% |
| **Hydrophobic Interaction** | F456-T27 | CR1 | 100% |
| | Y473-T27 | CR1 | 100% |
| | A475-T27 | CR1 | 100% |
| | F486-F28 | CR1 | 99% |
| | F486-L79 | CR1 | 99% |
| | F486-M82 | CR1 | 100% |
| | F486-Y83 | CR1 | 99% |
| | Y489-F28 | CR1 | 100% |
| | Y489-L79 | CR1 | 94% |
| | Y489-Y83 | CR1 | 99% |
| **Hydrogen Bonds** | Y449-D38 | CR3 | 23% |
| | N487-Y83 | CR1 | 65% |
| | Q493-E35 | CR2 | 49% |
| | Q498-Q42 | CR3 | 22% |
| | Q498-K353 | CR3 | 24% |
| | T500-Y41 | CR3 | 21% |
| | T500-D355 | CR3 | 51% |
| | Y505-E37 | CR3 | 31% |

**Table S2.** Pairwise interactions of $RBD_{BA.1}$ and $RBD_{BA.2}$ with ACE2.

| | Interaction | Location | $RBD_{BA.1}$-ACE2 | $RBD_{BA.2}$-ACE2 |
|---|---|---|---|---|
| **Salt bridges** | R403-E37 | CR2 | 69% | 33% |
| | K440-E329 | CR3 | 21% | 24% |
| | R493-E35 | CR2 | 100% | 99% |
| | R493-D38 | CR2 | 26% | 70% |
| | R498-D38 | CR3 | 59% | 29% |
| **Hydrophobic Interaction** | F456-T27 | CR1 | 100% | 100% |
| | Y473-T27 | CR1 | 100% | 100% |
| | A475-T27 | CR1 | 100% | 100% |
| | F486-F28 | CR1 | 98% | 98% |
| | F486-L79 | CR1 | 100% | 99% |
| | F486-M82 | CR1 | 99% | 99% |
| | F486-Y83 | CR1 | 98% | 98% |
| | Y489-F28 | CR1 | 100% | 100% |
| | Y489-L79 | CR1 | 90% | 90% |
| | Y489-Y83 | CR1 | 96% | 99% |
| | Y501-Y41 | CR3 | 100% | 100% |
| **Hydrogen Bonds** | Y453-H34 | CR2 | 16% | 19% |
| | A475-S19 | CR1 | 20% | 53% |
| | N477-S19 | CR1 | 21% | 55% |
| | N487-Y83 | CR1 | 46% | 75% |
| | T500-D355 | CR3 | 67% | 77% |
| | G502-K353 | CR3 | 74% | 83% |

**Table S3.** Predicted binding free energies using MMPBSA method

| Simulations | Binding Free Energy (kcal/mol) | Standard Deviation (kcal/mol) |
|---|---|---|
| $RBD_{WT}$ MD1 (PDB ID: 6M0J) | -29.5924 | 7.9485 |
| $RBD_{WT}$ MD2 (PDB ID: 6M0J) | -30.3688 | 6.6404 |
| $RBD_{BA.1}$ MD3 (PDB ID: 6M0J) | -40.3896 | 9.2167 |
| $RBD_{BA.1}$ MD4 (PDB ID: 6M0J) | -38.0986 | 10.2910 |
| $RBD_{BA.1}$ MD5 (PDB ID: 6M0J) | -36.1016 | 8.1722 |
| $RBD_{BA.1}$ MD6 (PDB ID: 7T9L) | -45.6993 | 8.3702 |
| $RBD_{BA.2}$ MD7 (PDB ID: 7T9L) | -46.9734 | 8.5512 |
| $RBD_{BA.2}$ MD8 (PDB ID: 7T9L) | -45.0240 | 9.6006 |
| $RBD_{BA.2}$ MD9 (PDB ID: 7T9L) | -43.7715 | 8.8759 |
| $RBD_{BA.2}$ MD10 (PDB ID: 7T9L) | -45.2938 | 9.3773 |

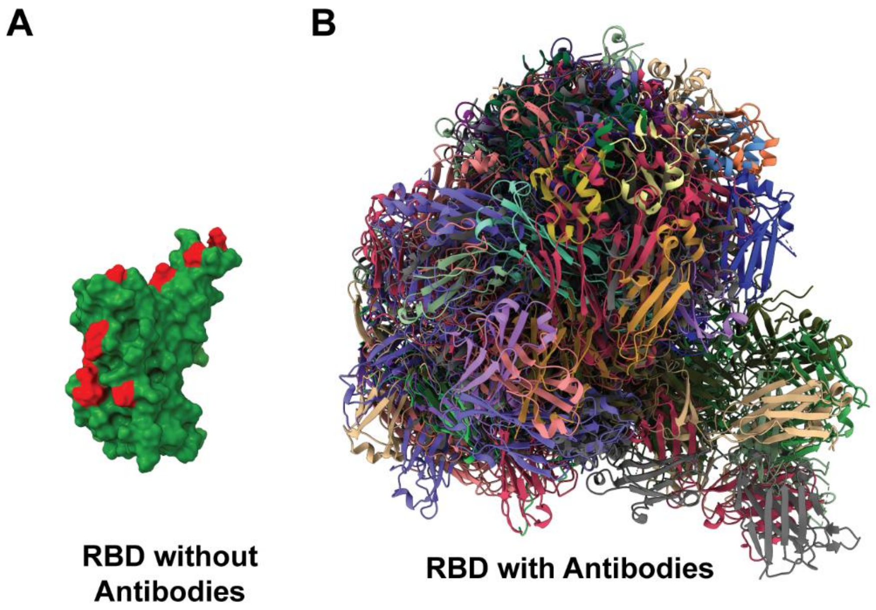


**Figure S1.** Antibody binding poses on RBD. (Left) Location of the RBD mutations of the Omicron variant (red). (Right) Structures of 160 antibodies were taken from the Protein Data Bank[1] and superimposed onto RBD.

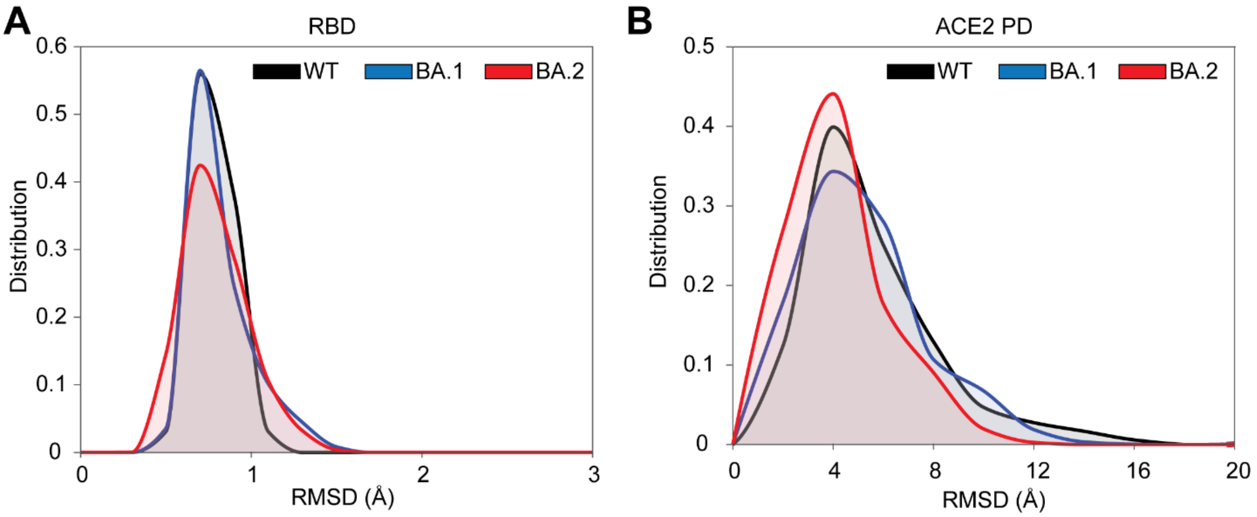


**Figure S2.** (A) Distribution of RMSDs of RBD from the $RBD_{WT}$-PD structure coordinates. (B) Distribution of RMSDs of PD from the $RBD_{WT}$-PD structure coordinates. From the RBD distributions representative RBD conformers showing the most probable RMSD values were selected. Using these conformers the RMSD between the following pairs $RBD_{WT}$:$RBD_{BA.1}$, $RBD_{WT}$:$RBD_{BA.2}$ and $RBD_{BA.1}$:$RBD_{BA.2}$ were evaluated as 0.75 Å, 0.82 Å, and 0.75 Å, hence showing minimal difference.
